# The *Azotobacter vinelandii* AlgU Regulon During Vegetative Growth and Encysting Conditions: A Proteomic Approach

**DOI:** 10.1101/2023.05.17.541114

**Authors:** Sangita Chowdhury-Paul, Iliana C. Martínez-Ortíz, Victoria Pando-Robles, Soledad Moreno, Guadalupe Espín, Enrique Merino, Cinthia Núñez

## Abstract

In the *Pseduomonadacea* family, the extracytoplasmic function sigma factor AlgU is crucial to withstand adverse conditions. *Azotobacter vinelandii,* a closed relative of *Pseudomonas aeruginosa*, has been a model for cellular differentiation in Gram-negative bacteria since it forms desiccation-resistant cysts. Previous work demonstrated the essential role of AlgU to withstand oxidative stress and on *A. vinelandii* differentiation, particularly for the positive control of alginate production. In this study, the AlgU regulon was dissected by a proteomic approach under vegetative growing conditions and upon encystment induction. Our results revealed several molecular targets that explained the requirement of this sigma factor during oxidative stress and extended its role in alginate production. Furthermore, we demonstrate that AlgU was necessary to produce alkyl resorcinols, a type of aromatic lipids that conform the cell membrane of the differentiated cell. AlgU was also found to positively regulate stress resistance proteins such as OsmC, LEA-1, or proteins involved in trehalose synthesis. A position-specific scoring-matrix (PSSM) was generated based on the consensus sequence recognized by AlgU in *P. aeruginosa*, which allowed the identification of direct AlgU targets in the *A. vinelandii* genome. This work further expands our knowledge about the function of the ECF sigma factor AlgU in *A. vinelandii* and contributes to explains its key regulatory role under adverse conditions.

## Introduction

*Azotobacter vinelandii,* a member of the *Psedomonadaceae* family, is a free-living bacterium having a strict aerobic metabolism with the particular capacity of fixing nitrogen in aerobiosis [1]. In contrast to *Pseudomonas* species, *Azotobacter* undergoes a differentiation process that culminates with the formation of cysts able to resist desiccation [2,3]. During *A. vinelandii* differentiation, several morphological and metabolic changes are observed, including the reduction in nitrogen fixation, the switch to anaerobic metabolism, or the loss of the flagella. It has also been reported that during this process, the bacterium replaces the phospholipids of the membrane by aromatic lipids, called alkylresorcinols (ARs) and alkyl pyrones [4], while granules of the polyester poly-hydroxy butyrate (PHB) accumulate in the central body as a reservoir of carbon and energy. The cell is surrounded by two layers, which are mainly composed of alginate, proteins, and ARs [2,3]; these layers are essential for the desiccation resistance of the mature cyst [5].

Several regulators have been identified to be essential for successful cyst formation in *A. vinelandii*. One of these regulators is the sigma factor AlgU, belonging to the family of extracytoplasmic function (ECF) sigma factors, and homolog to the stress response RpoE sigma factor of *Escherichia coli* [5,6]. These types of sigma factors induce gene expression in response to specific environmental stimuli [7]. The activity of ECF sigma factors is often controlled by a cytoplasmic membrane-bound anti-sigma factor that sequesters the ECF sigma factor [8]. Under inducing conditions, the ECF sigma factor is released, allowing it to interact with the RNA polymerase, activating gene expression. In *P. aeruginosa,* AlgU is normally sequestered by its cognate anti-sigma factor MucA. Proteolysis of MucA is initiated by the AlgW protease located in the periplasm, followed by cleavage by the MucP protease located in the inner membrane. MucA degradation is completed in the cytoplasm by AAA+ proteases such as ClpXP [9–16]. One of the signals triggering this proteolytic cascade is the accumulation of misfolded proteins in the periplasm. In *P. aeruginosa* AlgU directs RNA polymerase to activate the expression of a plethora of genes, including the alginate biosynthetic genes, encoded in an operon headed by *algD* [17].

In *A. vinelandii*, AlgU is also essential for expressing the alginate biosynthetic genes *algD* and *algC* [5;18–21]. In addition, AlgU was shown to be necessary for oxidative stress resistance [5] and for expression of *cydR* [22], encoding a repressor of *cydAB* genes (required for aerotolerant nitrogen fixation) and *flhDC*, which encode master regulators of flagella biogenesis [22].

To elucidate the global molecular changes that occur during encystment, in a previous work we determined the proteome of the *A. vinelandii* cells undergoing differentiation [23]. We identified proteins differentially expressed during encystment induction with respect to vegetative cells. We found modifications in the abundance of proteins involved in nitrogen fixation, flagella synthesis and cell division, which agrees with the biochemical, morphological and physiological changes known to take place during this process [23].

In the current work, proteins under the control of the sigma factor AlgU in vegetative or encysting conditions were identified using a proteomic approach. They include proteins for processes previously known to be dependent on AlgU, but our study also revealed that AlgU was necessary to produce ARs and trehalose or to positively regulate stress resistance proteins such as OsmC, MdaB, and LEA-1. A position-specific scoring matrix (PSSM) was generated based on the consensus sequence recognized by AlgU in *P. aeruginosa*, allowing it to confirm some of the genes positively regulated by AlgU in *A. vinelandii*.

## Materials and methods

### Strains and cultivation conditions

The *A. vinelandii* wild-type AEIV strain (also named E strain) [24] and its *algU* or *mucA* derivative mutants, named AEalgU and AEA8 [5, 21], respectively, were used in this study. *A. vinelandii* cells were routinely cultivated in Burk’s medium; sucrose (20 gL^−1^) was used as a carbon source (Burk’s-sucrose medium) [1]. The composition of the culture medium has been previously reported [25].

For encystment induction, the *A. vinelandii* wild-type strain and its *algU* derivative mutant AEalgU, was cultured for 48 h in Burk’s-sucrose medium. The cells were harvested and washed three times with Burk’s solution (Burk’s medium without carbon source). The cells were then resuspended in fresh medium supplemented with 0.2% v/v of *n*-butanol as the sole carbon source and incubated for 48 h at 30°C.

### Analytical methods

Protein concentration was quantitated as described previously [26]. The activity of β-glucuronidase in cells grown in liquid medium was determined as reported before [27] with some modifications; *A. vinelandii* cells were permeabilized using 0.013%(W/V) lysozyme and incubating at 37 °C/5 min, followed by the addition of 0.13 % (V/V) Triton before the enzymatic assay. One U corresponds to 1 nmol of O-nitrophenyl-β-D-glucuronide hydrolyzed per minute per μg of protein. Quantification of ARs and ARs qualitative visualization on agar plates was conducted using the Fast Blue colorimetric assay [28]. Details of the adapted methods for *A. vinelandii* are described elsewhere [29]. All experiments were conducted in triplicates; the results presented are the averages of the independent runs. Statistical analysis was carried out using a Student’s t-test (p = 0.05).

### Construction of an *algC-gusA* transcriptional fusion

A fragment of 306 bp, spanning a region from nucleotides −342 to −36 with respect to the ATG translation initiation codon of *algC*, was PCR amplified using oligonucleotides algC-gusF and algC-gusR (S1 Table). This fragment was subsequently cloned into plasmid pUMATcgusAT [30] as an XbaI-EcoRI fragment (the restriction sites were included in the corresponding oligonucleotides), thus generating plasmid pCN62. Strain AEIV was transformed with plasmid pCN62, previously linearized with NdeI endonuclease, and transformants Tc^r^ were selected. The strain generated, carrying an *algC-gusA* transcriptional fusion, was named CNL35.

### Cell fractionation and protein sample preparation

The proteomic analysis was conducted using soluble protein extracts derived from three independent cultures (biological replicates) from the wild-type strain and from mutant *algU*, grown under vegetative and encysting conditions. Then, the AlgU regulon was obtained by comparing the proteins expressed in the *algU* mutant with those expressed in the wild-type strain in both conditions, vegetative and encystment [23]. Cells from vegetative (24 h/Burk’s-sucrose medium) and encysting conditions (48 h/ Burk’s-butanol medium) were harvested by centrifugation at 1900 x g for 10 min at 4°C, washed, and resuspended in 10mM of sodium-phosphate buffer of pH 7.4. Cell fractionation and sample preparation were conducted as described [23].

### LC-MS/MS analysis and identification/quantification of proteins

A detailed description of the analysis and protein quantification procedures has been previously reported [23]. LC-MS/MS analysis was performed at the IRCM Proteomics Discovery Platform of the Montreal Clinical Research Institute as described previously. Raw files obtained from Orbitrap Q-Exactive spectrometer were acquired using Mascot 2.3 (Matrix Science) against a database of *A. vinelandii* DJ strain from NCBI (taxon identifier 322710). Data analysis was performed using Scaffold (http://www.proteomesoftware.com/products/scaffold/download/) [31]. Protein expression in the *algU* mutant was considered significantly different only if protein ratios differed more than two-fold with respect to the expression observed in the wild-type strain AEIV. The Student’s t-test was also done for each sample set having three different biological replicas and a threshold level of 0.05 was considered for selecting the proteins.

### Protein functional classification analysis

Protein functional classification and KEGG pathways analysis of differentially modulated proteins was conducted using the Kyoto Encyclopedia of Genes and Genomes database (www.genome.jp/kegg) [32]. The String 10 database was used to determine protein–protein interaction network [33].

### Quantitative analysis of mRNA levels

Strains AEIV, AEalgU and AEA8 were cultured in Burk’s-sucrose or in Burk’s-butanol medium for 24 or 48 h, respectively. Cells were collected by centrifugation, and the total RNA was extracted as described [34]. Details of DNA contamination removal, cDNA synthesis and qPCR amplification conditions are reported elsewhere [25]. qPCR assays were performed with a Light Cycler 480 II instrument (Roche), using the Maxima TM SYBR Green/ROX qPCR Master Mix (2X) kit (Thermo Scientific). The sequences of the primer pairs used are listed in S1 Table. Three biological replicates (independent cell cultures) were performed, with three technical replicates for each one. Similar results were obtained for the transcription of all measured genes in the repetitions. Relative mRNA transcript levels were determined in relation to *gyrA* (Avin15810) mRNA, as reported previously [25]. A non-template control of each reaction was included for each gene. The quantification technique used to analyze the generated data was the 2^−Δ,ΔCT^ method reported previously [35].

### Identification of orthologous genes

Orthologous genes were defined using “bidirectional best hits” criteria where reciprocal best hits were identified by pairwise comparisons using BLAST (BLASTP version 2.12.0+) [36]. The results were filtered using a cut off E-value of 1×10^5^ and a query and subject coverage of at least 50%.

### Operon predictions

Operon predictions were performed using the method previously reported [37], which is based on the intergenic distance between codirectional transcribed genes, and the functional relationship of their protein products defined in the STRING database [38].

### *In silico* prediction of AlgU-dependent promoters

Experimentally determined AlgU-dependent promoters in *P. aeruginosa* (S2 Table) were used as a reference to construct a Position-Specific Scoring Matrix (PSSM) for the *de novo* motif detection using *ad hoc* developed PERL program [39–47]. Our PSSM was based on the tight consensus sequence [(−35) **GAA**CTT-N16/17-(−10)**TC**tgA (highly conserved residues in bold capital letters; conserved residues in capital letters)] reported for most of the AlgU-dependent promoters. Our PSSM used only three different values: 1, 0.9, and 0.81 associated with invariant, well-conserved, and conserved nucleotides within the promoter sequences, respectively. We only considered as likely promoters those whose −10 and −35 consensus boxes were separated by 16 or 17 nt. For our AlgU-dependent promoter search, we scanned the first 250 nt upstream of every gene regardless of the gene position within their corresponding operons. This consideration allowed us to identify internal promoter sequences within coding regions. The promoter score was obtained by multiplying the values associated with the nucleotides of the region analysed, based on our PSSM in such a way that sequences with 100% conserved nucleotides obtained a score of 1, sequences with a substitution of one nucleotide in one position obtained a score of 0.9 and sequences with one substitution in a highly conserved region or two substitutions in conserved regions, obtained a score of 0.81. The search was performed for the *P. aeruginosa* PAO1 (tax id: 208964), *A. vinelandii* DJ (tax id: 322710), *Azotobacter chroococcum* Ac-8003 (tax id: 1328314) and *Pseudomonas fluorescens* SBW25 (tax id: 216595) genomes. The latter two served as comparative genomes for AlgU-predicted promoters in *A. vinelandii* and *P. aeruginosa,* respectively.

## Results and discussion

### The activity of AlgU increases upon encystment induction

The essential role of the sigma factor AlgU during *A. vinelandii* encystment has been clearly demonstrated in previous works from our laboratory [5,22]. As the activity of this type of sigma factor is highly regulated at the protein level, we reasoned that such activity would be increased upon encystment induction. To explore this assumption, we constructed a P*algC*-*gusA* transcriptional fusion since expression of *algC* was previously shown to be under the direct control of AlgU [19]. The activity of AlgU was estimated during vegetative growth in Burk’s-sucrose liquid medium and during encysting conditions, induced with 0.2% *n*-butanol. As a negative control, this transcriptional P*algC*-*gusA* fusion was tested in an *algU^−^* genetic background. In vegetative conditions, at 24 h, the activity of AlgU reached the highest point. Thereafter the activity slightly decreased at 48 h (Fig 1A). As expected, we detected a basal level of P*algC* activity in the negative control that did not significantly change along the growth curve. 24 h after induction of encystment, the activity of AlgU doubled and remained high for the following 24 h; then, it gradually dropped to levels like those observed at the end of vegetative growth (Fig 1B). Therefore, samples from 24 h for the vegetative condition and from 48 h for the encystment condition, were used for further analysis of the AlgU regulon.

**Fig 1.**
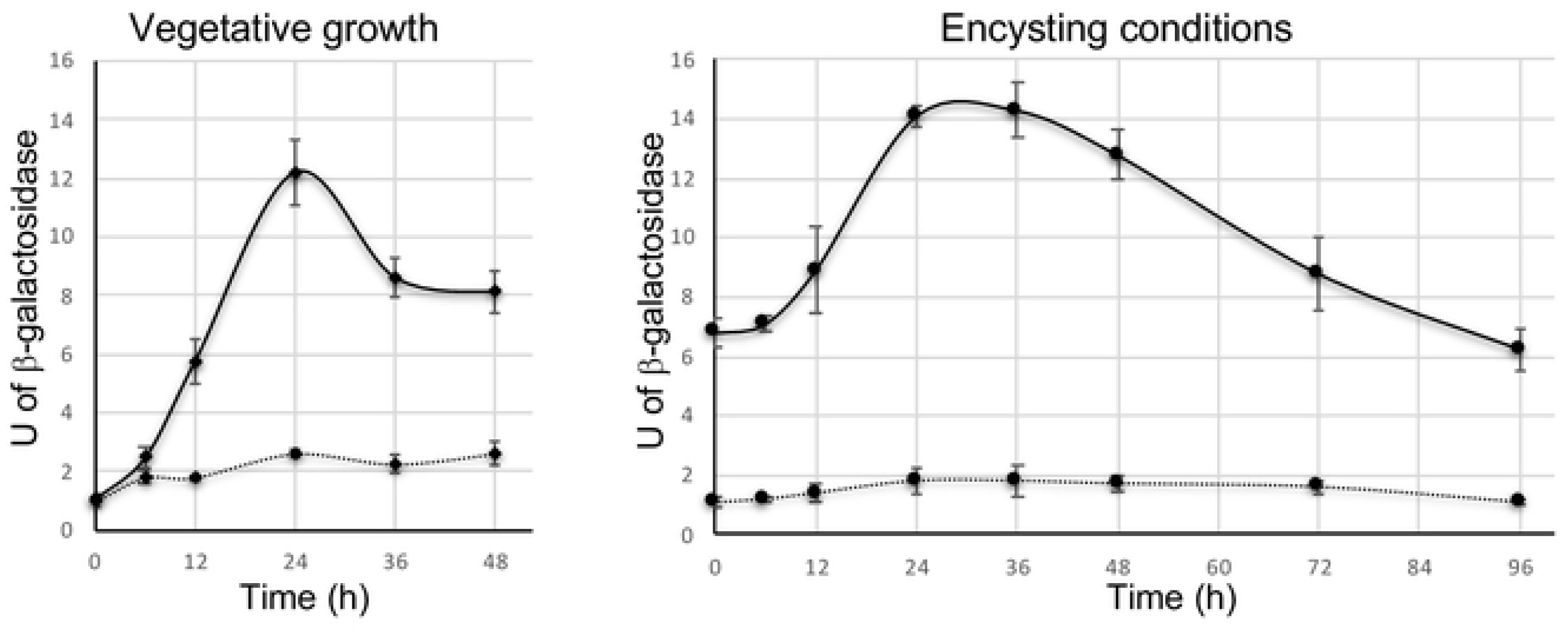
Activity of the AlgU sigma factor in *A. vinelandii*. A transcriptional *PalgC-gusA* fusion was used to estimate the activity of AlgU in the background of the wild-type strain AEIV (solid line) or in the *algU* mutant (dotted line). Cells were cultured in minimum Burk’s medium supplemented with sucrose (vegetative growth) or *n*-butanol (encystment-inducing conditions) as the sole carbon source for the indicated time. The bars of standard deviation from three independent experiments are shown.

### The AlgU proteome during vegetative growth

To compare the total proteomic profile in cytoplasmic fraction of the *algU* mutant, we have analyzed the protein samples through Hybrid Quadrupole-Orbitrap Mass Spectrometer, which combines quadruple precursor ion selection with high-resolution, accurate-mass (HRAM) Orbitrap detection. The total expressed proteins in the absence of AlgU (S3 Table) were compared to the previously reported proteins present in the wild-type strain AEIV [23]. After evaluating the data generated from orbitrap, 126 proteins were found to be differentially expressed in the *algU* mutant, among which, 50 proteins were downregulated (S4 Table), and 76 proteins were upregulated (S5 Table), after 24 hours of vegetative growth, as compared to the wild-type strain.

With the aim of understanding the roles of the differentially expressed proteins due to the absence of AlgU under vegetative conditions, we mapped the genes encoding the differentially expressed proteins to their corresponding terms in the KEGG database. We identified 20 main pathways for the 126 differentially expressed proteins (Fig 2A). Besides the group of proteins of unknown function, the most represented groups corresponded to energy, carbohydrate and amino acid metabolism. Protein networks generated by String 10 software, identified 81 protein interactions (for a confidence interaction score of 0.7), for the 126 proteins. Interaction nodes include those for proteins related to central and lipid metabolism, flagella biogenesis, trehalose synthesis, among others (S1 Fig).

**Fig 2.**
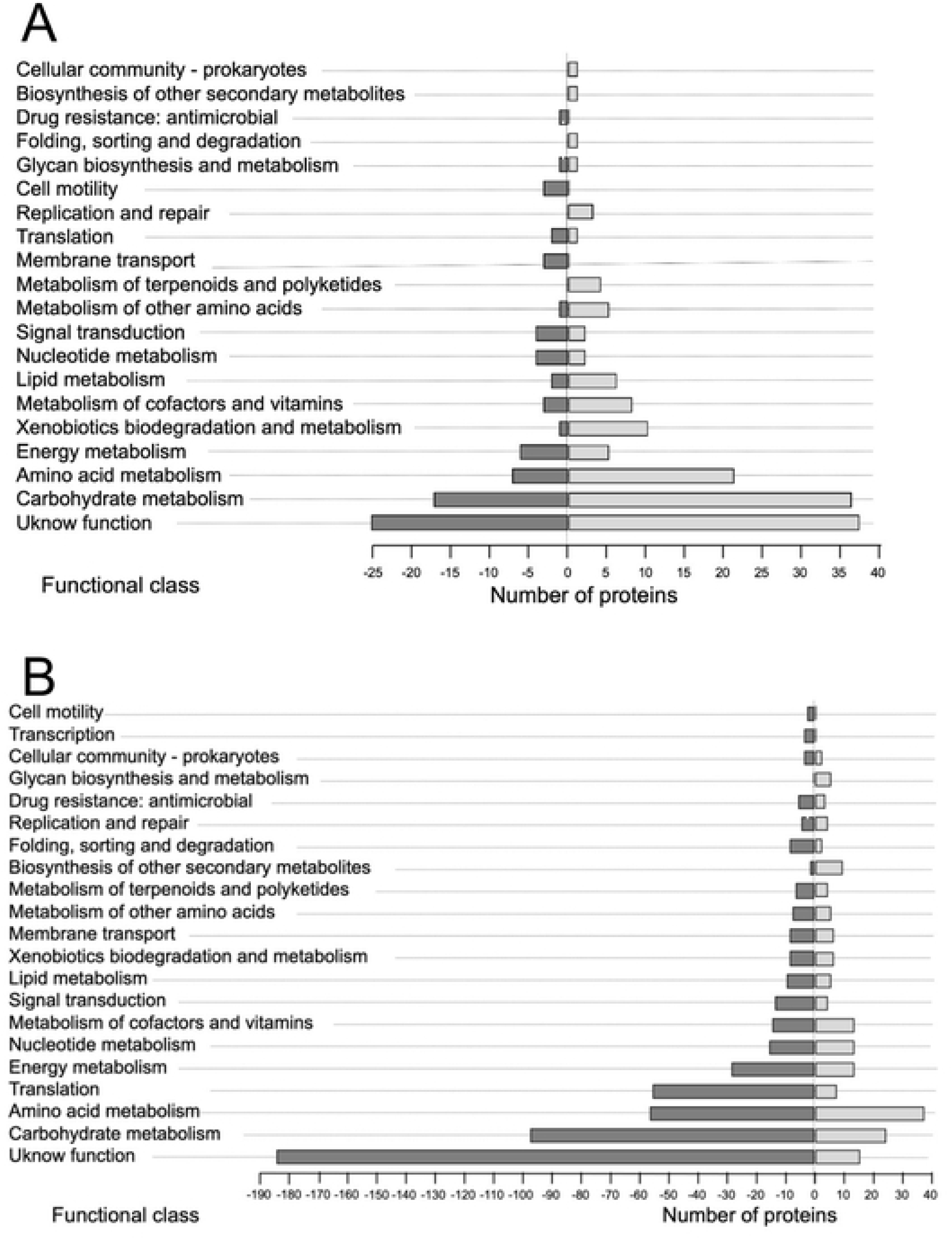
KEGG pathway enrichment of proteins of the AlgU regulon in *A. vinelandii*. The analysis was conducted for the differentially expressed proteins in the absence of the sigma factor AlgU under vegetative (A) or encystment-induced (B) conditions. Positive and negative axes represent the numbers of up- or downregulated proteins, respectively.

### Analysis of proteins under the control of AlgU during vegetative conditions

The abundance of the corresponding mRNAs for 6 proteins found under the control of AlgU was evaluated by qPCR. We reasoned that accumulation of such mRNAs would be diminished in the *algU* mutant but would be increased in a mutant lacking the anti-sigma factor MucA, thus showing elevated AlgU activity. The results are presented in Table 1.

**Table 1.**
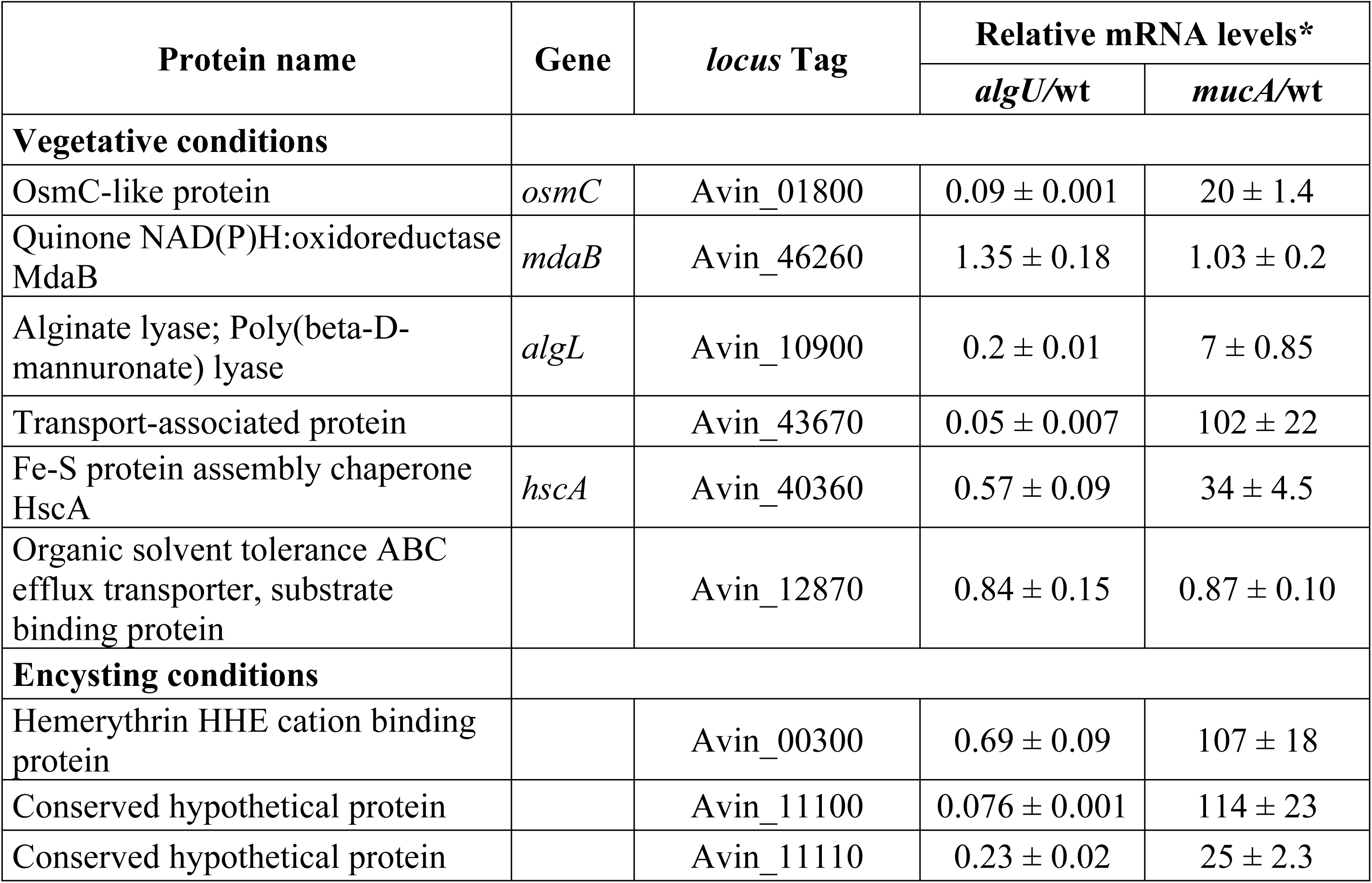
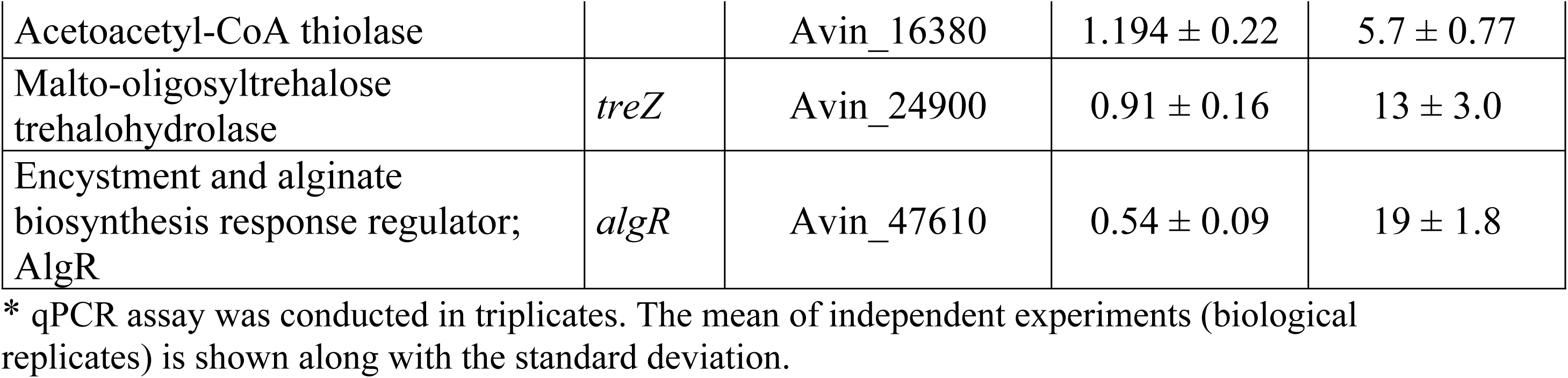
Relative mRNA levels of some genes encoding proteins under the positive control of AlgU.

| Protein name | Gene | locus Tag | Relative mRNA levels* |  |
| --- | --- | --- | --- | --- |
|  |  |  | *algU*/wt | *mucA*/wt |
| Vegetative conditions |  |  |  |  |
| OsmC-like protein | *osmC* | Avin_01800 | 0.09 ± 0.001 | 20 ± 1.4 |
| Quinone NAD(P)H:oxidoreductase MdaB | *mdaB* | Avin_46260 | 1.35 ± 0.18 | 1.03 ± 0.2 |
| Alginate lyase; Poly(beta-D-mannuronate) lyase | *algL* | Avin_10900 | 0.2 ± 0.01 | 7 ± 0.85 |
| Transport-associated protein |  | Avin_43670 | 0.05 ± 0.007 | 102 ± 22 |
| Fe-S protein assembly chaperone HscA | *hscA* | Avin_40360 | 0.57 ± 0.09 | 34 ± 4.5 |
| Organic solvent tolerance ABC efflux transporter, substrate binding protein |  | Avin_12870 | 0.84 ± 0.15 | 0.87 ± 0.10 |
| Encysting conditions |  |  |  |  |
| Hemerythrin HHE cation binding protein |  | Avin_00300 | 0.69 ± 0.09 | 107 ± 18 |
| Conserved hypothetical protein |  | Avin_11100 | 0.076 ± 0.001 | 114 ± 23 |
| Conserved hypothetical protein |  | Avin_11110 | 0.23 ± 0.02 | 25 ± 2.3 |
| Acetoacetyl-CoA thiolase |  | Avin_16380 | 1.194 ± 0.22 | 5.7 ± 0.77 |
| Malto-oligosyltrehalose trehalohydrolase | *treZ* | Avin_24900 | 0.91 ± 0.16 | 13 ± 3.0 |
| Encystment and alginate biosynthesis response regulator; AlgR | *algR* | Avin_47610 | 0.54 ± 0.09 | 19 ± 1.8 |

\* qPCR assay was conducted in triplicates. The mean of independent experiments (biological replicates) is shown along with the standard deviation.

As our key target was to study the proteins under AlgU regulation, we were very interested in the proteins which are most affected (missing) due to the *algU* mutation. We have found 18 proteins that completely disappeared due to the mutation in *algU* during vegetative growth as compared to wild type (Table 2 and S4 Table).

**Table 2.**
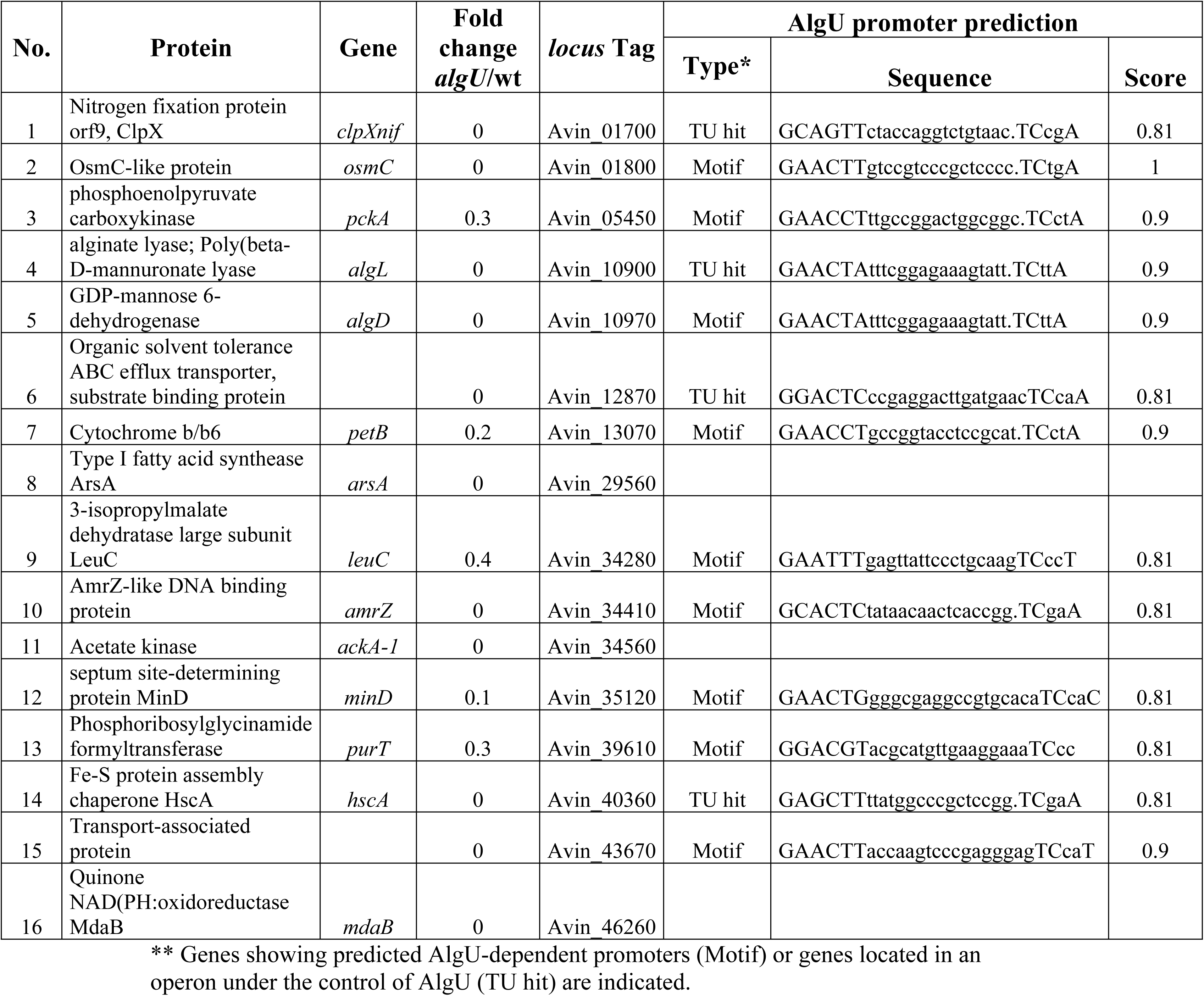
Selected proteins under the positive control of AlgU under vegetative growing conditions

As anticipated, some proteins for alginate biosynthesis were found to be down-affected in the *algU* mutant, including the GDP-mannose 6-dehydrogenase (AlgD), which is the enzyme catalyzing the key step in alginate biosynthesis or AlgL, an alginate lyase that also serves as part of the multi-protein alginate-secretion complex [48–50]. As expected, the abundance of the *algL* mRNA in the *algU* mutant was 5-fold reduced, but it was increased 7-fold in the *mucA* genetic background, as revealed by qPCR (Table 1).

Besides these, the OsmC-like protein was found to be absent in the *algU* mutant. Osmotically inducible protein C (OsmC) is a protein found in *E. coli* during stress conditions such as salt stress [51] or in a medium of low osmotic pressure [52]. In oxidative stress OsmC-induced cells were found to be highly viable whereas an *osmC* mutant showed more sensitivity to butanol in the exponential growth phase and to H_2_O_2_ and butanol in the stationary phase [53], indicating that OsmC may play a role as a scavenger for specific ROS [51]. In *P. aeruginosa* OsmC is part of the AlgU regulon [41]. In *A. vinelandii* we have shown that expression of OsmC is completely dependent on AlgU during both, vegetative and encysting conditions (see below). Accordingly, the *osmC* mRNA was 10-fold reduced in the *algU* mutant, but 20-fold higher in the *mucA* mutant (Table 1). As in *P. aeruginosa*, transcription of *osmC* seems to be under direct regulation of AlgU as we found a potential AlgU promoter in its regulatory region (Table 2). Description of AlgU promoters’ prediction in the genome of *A. vinelandii* is detailed in the last section of Results.

The quinone oxidoreductase MdaB protein was absent in the *algU* mutant. This protein was first identified as a modulator of drug activity in *E. coli* [54]. Quinone oxidoreductases were shown to reduce quinone substrates via an intensive two-electron mechanism which play key roles to maintain a pool of reduced quinols that contribute to antioxidant defense in *E. coli* and *P. aeruginosa* [55–57]. MdaB of *P. aeruginosa* shares 70% identity with its orthologous in *A. vinelandii*. Accumulation of the *mdaB* mRNA was not diminished in the *algU* mutant (Table 1), nor affected in the *mucA* mutant implying that the effect of AlgU on the expression of MdaB might be post-transcriptionally.

An iron-sulfur protein assembly chaperone HscA was undetected in the *algU* mutant. HscA is an Hsp70 class molecular chaperone previously described in many bacteria, including *E. coli*, *P. aeruginosa* and *A. vinelandii* [58–61]. In *A. vinelandii* the *hscA* gene is a component of the *isc* operon which is responsible for Fe-S cluster biogenesis and helps in the maturation of [2Fe–2S] proteins [58]. Several genes of the *isc* operon, including *hscA*, were found to be essential for vegetative growth in the OP genetic background. The OP strain is a naturally occurring *algU* mutant. As we detected no HscA protein in the *algU* mutant derived from strain AEIV, it is likely to propose a basal expression level of the *isc* operon undetectable by our proteomic approach. Indeed, qPCR assay showed that in the *algU* mutant, the levels of the *hscA* mRNA were diminished by about 50%, but its accumulation increased 34-fold in the background of mutant *mucA,* indicating a positive dependence on AlgU (Table 1). This agrees with a predicted AlgU promoter directing the transcription of the *hcsA* containing operon (Table 2). Another important function of this operon is to defend against oxidative stress [61–64]. All the above three proteins (OsmC, MdaB and HscA) related to counteracting the stress response, and found under the control of AlgU, explain the previous observation of the reduced survival of an *algU* mutant in oxidative stress conditions [5].

Another protein absent in the *algU* mutant is an ABC efflux transporter protein (Avin12870). ATP binding cassette (ABC) efflux transporters proteins use energy to remove solvents from the cells to the outer medium. ABC transporters translocate a wide variety of substrates, including amino acids, peptides, ions, sugars, toxins, lipids and drugs in several bacteria including *Pseudomonas* and *E. coli* [65]. These proteins discharge the toxic compounds from the cell to the external medium which is a relevant mechanism in the solvent-tolerance of bacteria [66]. Of note, differences in accumulation of the Avin_12870 mRNA was not detected in either, the *algU* or the *mucA* mutant with respect to the WT strain (Table 1), even though an AlgU dependent promoter was detected for its transcriptional unit.

Our analysis also revealed a transport associated protein (Avin_43670) which has a BON domain. This domain was found in an OsmY protein of *E. coli,* that was reported to protect the cell against stress, especially during osmotic shock by contacting the phospholipid interfaces surrounding the periplasmic space [67]. In *A. vinelandii* the Avin_43670 transport-associated protein may also help to perform these functions by binding or interacting with the phospholipid membrane. The Avin_43670 gene shows a putative AlgU promoter implying a direct transcriptional regulation. In agreement with this, accumulation of the corresponding mRNA was diminished in the *algU* mutant, whereas it showed a strong upregulation, of about 102-fold in the *mucA* mutant.

### The *A. vinelandii* AlgU regulon during encystment

As described in Materials and Methods, the proteome of AlgU was also determined at 48 h of encystment induction. The total expressed proteins in the *algU* mutant were obtained (S6 Table), and their abundance was compared to those of the wild-type strain [23]. We found 305 proteins downregulated (S7 Table) and 184 ones upregulated (S8 Table) in the *algU* genetic background. The vast number of proteins whose expression was affected in the absence of AlgU might reflect the central role of this sigma factor during *A. vinelandii* encystment.

Based on the KEGG database, the differentially expressed proteins due to the *algU* mutation are involved in several functions, including carbohydrate and amino acid metabolism, translation and signal transduction, among others (Fig 2B).

Protein interaction networks were generated by String 10 software (confidence cutoff of 0.7), for proteins downregulated (S2 Fig) or upregulated (S3 Fig) in the absence of AlgU. The number of interaction nodes for both data sets was significantly higher than expected, revealing a connection among the identified proteins. This analysis also confirmed the positive role of AlgU in oxidative phosphorylation, amino acid metabolism, synthesis of glycogen and trehalose or alginate production, among others, as revealed by the cluster of identified functional groups. Furthermore, a negative role of AlgU in amino acids and aminoacyl-tRNA biosynthesis was also identified (S2 and S3 Fig).

### Analysis of proteins under the control of AlgU during encystment-induced conditions

After 48 h of encystment induction, 183 proteins were not detected in the *algU* mutant when compared to the wild-type strain (S7 table). Strikingly, 50 of such proteins were strongly expressed during encystment in the wild-type strain, as compared to its own vegetative growth (Table 3 and S7 Table) [23], revealing AlgU targets specifically associated to this differentiation process. The expression levels of six of these proteins were also examined by qPCR (Table 1) and the results are discussed below.

**Table 3.**
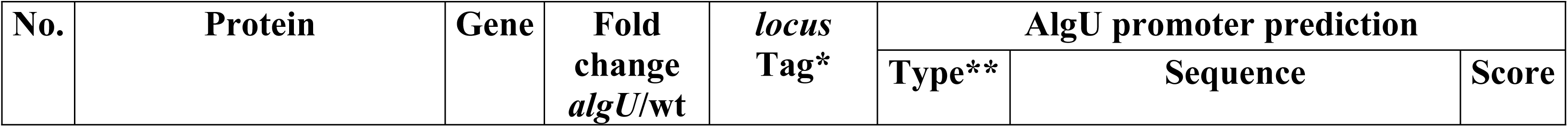
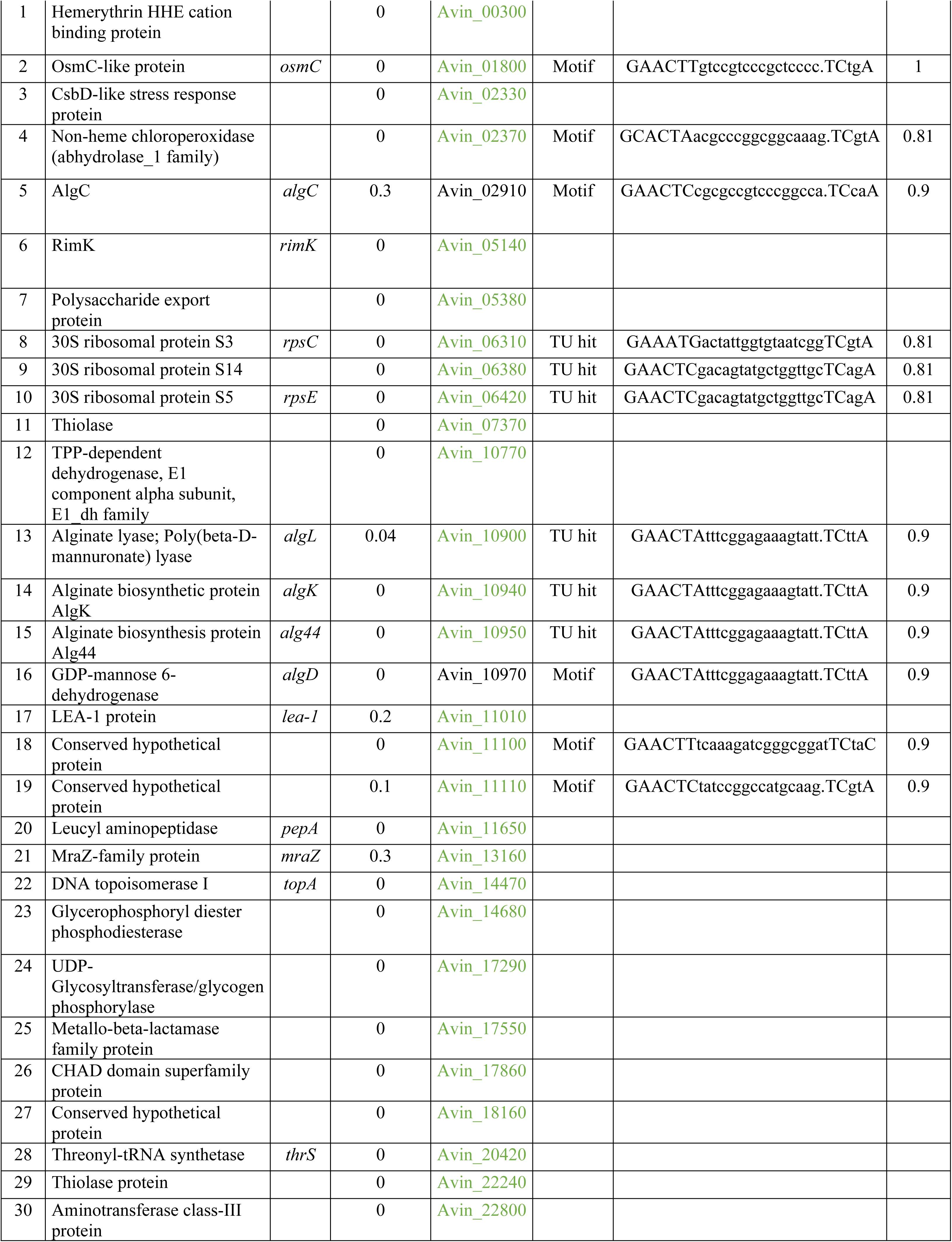
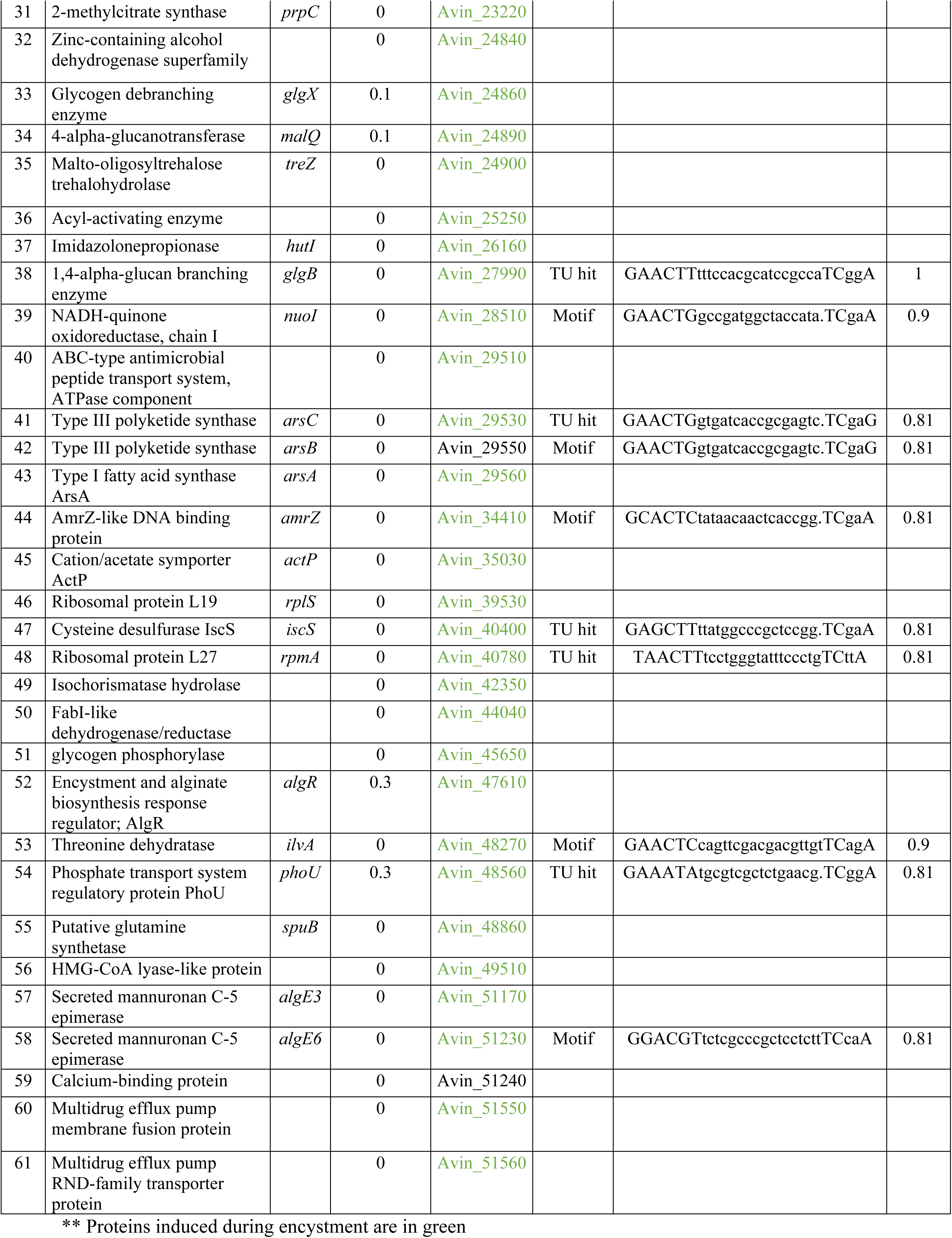
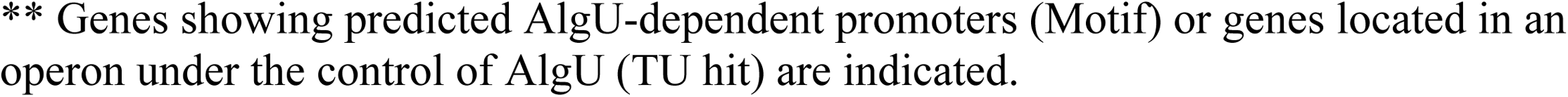
Selected proteins positively control by AlgU during encysting conditions

The abundance of several alginate biosynthetic proteins was reduced in the *algU* mutant, confirming the positive role of AlgU for alginate production during cyst formation. AlgD, AlgE3 and AlgE6 (Secreted mannuronan C-5 epimerase) were totally missing in the *algU* mutant along with ORF Avin51240, encoding a calcium-binding protein important for the activity of the mannuronan C-5 alginate epimerases. AlgL and AlgC were 0.05 and 0.3-fold downregulated as compared to wild type, respectively. The response regulator AlgR, which is essential for cyst formation [68], was also found to be 0.4-fold downregulated in mutant *algU*. The AlgU-dependent transcription of *algR* was further demonstrated by qPCR as *algR* mRNA levels were reduced in the *algU* mutant but were enhanced 19-fold in the *mucA* mutant (Table 1). In *P. aeruginosa* AlgU recognizes the promoter of *algR* for transcription initiation [42,43]. In *A. vinelandii*, however, the positive effect of AlgU on *algR* might be indirect as the *algR* promoter does not show consensus sequences recognized by this sigma factor (Table 3) [68].

The hemerythrin HHE cation binding protein (Avin_00300), strongly expressed during encystment of the wild-type strain [23,30], was undetectable in the *algU* mutant. This effect seems to be at the transcriptional level as the amount of the corresponding mRNA was reduced in the *algU* mutant, but it was 107-fold higher in the *mucA* mutant (Table 1). Hemerythrin is a non-heme, iron-containing protein that binds to oxygen [69]; it has been proposed to regulate interactions between cellular enzymes and oxygen or to be involved in the transport of oxygen within the cell [70,71]. However, its exact role in *A. vinelandii* differentiation remains unknown.

Other proteins involved in synthesizing trehalose such as GlgX, MalQ and TreZ (Avin_24860, Avin_24890 and Avin_24900, respectively), and previously shown to be induced during encystment, were also downregulated in the *algU* mutant. The control of these genes by AlgU might be complex, involving different layers of regulation, as the levels of *treZ* mRNA was not significantly reduced in the absence of this sigma factor, but it was 13-fold higher in the *mucA* genetic background. Late embryogenesis abundant (LEA) proteins conform a large family associated with resistance to abiotic stress. *A. vinelandii* cysts expresses LEA-1 protein essential for the survival of the differentiated cell in dry conditions and high temperatures [72]. Expression of LEA-1 was AlgU dependent, as its accumulation was 5-fold reduced in the *algU* mutant when compared to the wild-type strain.

In several *Pseudomonas* spp. and in *E. coli*, the protein RimK is involved in modifying the ribosomal protein RpsF, affecting the translation of several key genes necessary to survive adverse conditions [73,74]. In *A. vinelandii* RimK was previously shown to be strongly induced during encystment [23]. Our proteomics results indicate that its expression depends on AlgU as in its absence, RimK (Avin_05140) was undetectable. The reported role of RimK in controlling the activity of ribosomes may be related to the strong downregulation of multiple 50S and 30S ribosomal proteins in the absence of AlgU, highlighting the role of this sigma factor on protein translation (S7 Table and S2 Fig).

It is worth mentioning that the regulon detected for AlgU during encystment contains a total of 49 hypothetical proteins, which implies the existence of additional cellular processes yet to be defined under the control of this sigma factor. Two hypothetical proteins (Avin_11100 and Avin_ 11110), upregulated during encystment, were missing in the *algU* mutant. This effect seems to be at the transcriptional level as the amount of the corresponding mRNAs was negligible in the *algU* mutant, but it was strongly enhanced in the *mucA* genetic background (Table 1). This result agrees with the presence of an AlgU-dependent promoter in their regulatory regions (Table 3).

### AlgU controls ARs production

Upon encystment, 95% of the cell membrane phospholipids are replaced by the phenolic lipids alkyl-resorcinols (ARs) and alkyl-pyrones [4]. Interestingly, expression of proteins ArsA, ArsB and ArsC for the synthesis of these lipids was totally suppressed in the *algU* mutant, implying that cell membrane phospholipids replacement does not occur in the absence of AlgU (Table 3 and S7 Table). Indeed, ARs in encystment-induced cells were not detected in the *algU* mutant, in contrast to the wild-type strain (Fig 3A). Accordingly, the synthesis of fatty acids seems to remain active in the *algU* mutant, as suggested by the up regulation of some proteins involved in this pathway (Avin_14930, Avin_15000, Avin_29050 and Avin_44250) (S8 Table and S3 Fig).

**Fig 3.**
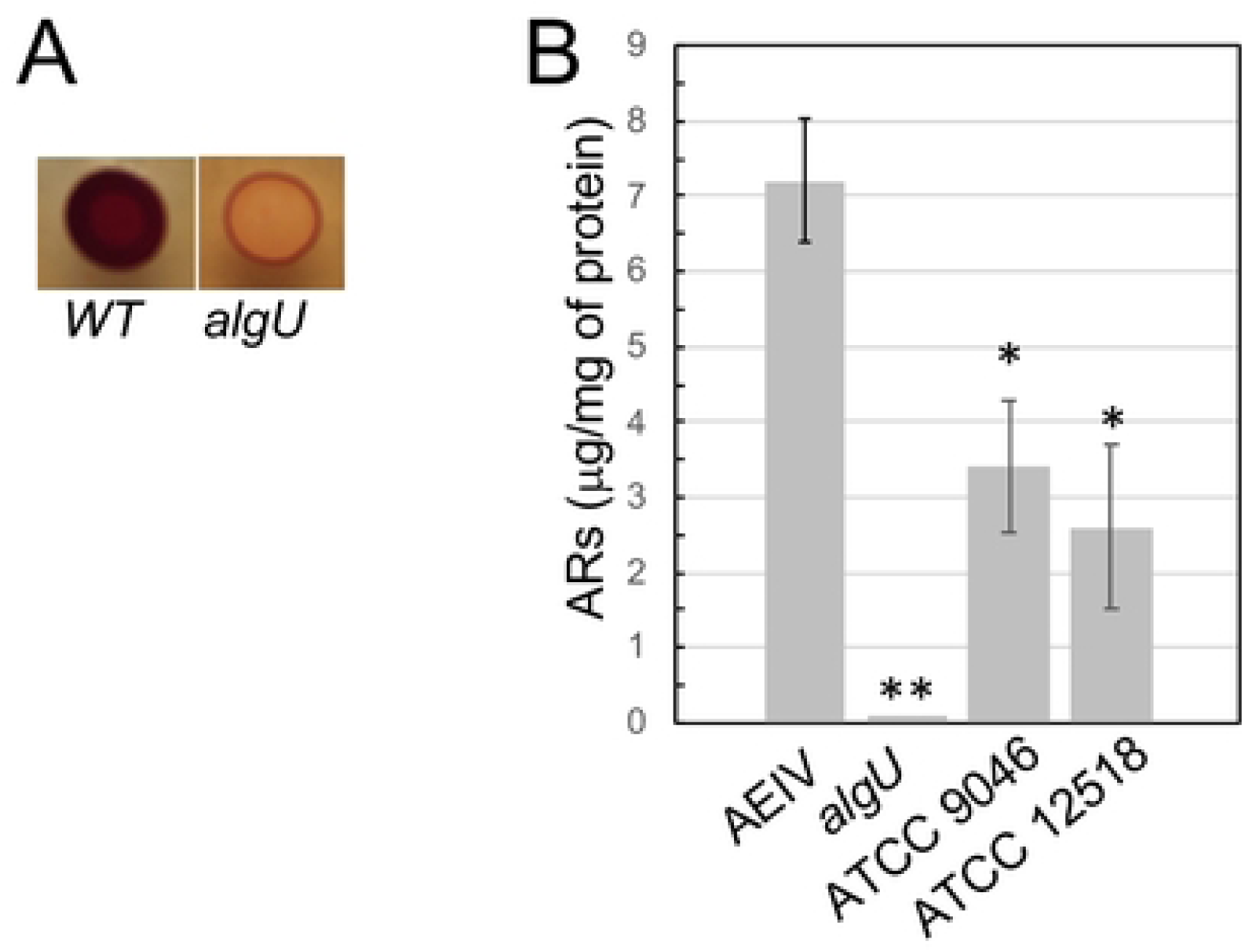
The production of ARs is impaired in the absence of AlgU. A. Staining with Fast Blue B of ARs produced by strain AEIV (WT) and by its derivative *algU* mutant under encysting conditions. Cells were grown on Burk’s-butanol for 48 h before staining. B. ARs quantification under vegetative growth. The wild-type strains AEIV, ATCC 9046 and ATCC 12518, and the AEIV derivative mutant *algU* were cultivated in Burk’s-sucrose medium for 24 h prior to ARs extraction. The bars of standard deviation from three independent experiments are shown. The asterisks denote statistical significance (unpaired Student’s *t*-test, *P<0.05; **P<0.01) when compared to the wild-type strain AEIV.

Previous reports indicated the production of ARs in glucose aging cultures of *A. vinelandii* strain ATCC12837 [75]. Similarly, quantification of ARs production by the wild-type strain AEIV at 24 h of vegetative growth, confirmed the presence of these phenolic lipids (Fig 3B). The same was true for *A. vinelandii* wild-type strains ATCC 9046 and ATCC 12518. Of note, these two strains produced ARs in a lower amount when compared to strain AEIV. This result agrees with our proteomic data indicating the expression of ArsA in WT vegetative cells. The regulation of ARs by AlgU also occurs during vegetative growing conditions; ArsA was undetectable in mutant *algU* (Table 2) and correlated with the absence of these lipids in this genetic background (Fig 3B). Furthermore, q-PCR suggested that the positive effect of AlgU on ARs production occurs at the transcriptional level since mRNA accumulation of *arsA* and *arsB* was abrogated in the absence of AlgU under both, vegetative or encystment-induced conditions (0.0015 ± 1×10^−4^ and 0.0008 ± 1.6×10^−5^ for *arsA* and *arsB*, respectively), when compared to the wild-type strain.

### Identification of AlgU binding motifs

To identify potential targets directly regulated by AlgU in *A. vinelandii*, AlgU-dependent promoters were predicted based on the consensus sequence recognized by this sigma factor in *P. aeruginosa*. *A. vinelandii* and *P. aeruginosa* are phylogenetically related and their AlgU binding motifs, so far reported, are very well conserved [6,19,76].

A Position-Specific Scoring Matrix (PSSM), was developed as described in Materials and methods, using as a reference experimentally determined *P. aeruginosa* AlgU promoters previously reported (i.e., by mapping the 5’ end of the corresponding mRNA) (S2 Table) [39–47]. This PSSM was used to search the genome of *P. aeruginosa* PAO1. A total of 134 AlgU promoters with a score ≥ 0.9 were predicted (motifs showing invariant or well-conserved consensus sequences), and 486 with a score of 0.81 (motifs showing conserved consensus sequences) (S9 Table).

The *P. aeruginosa* AlgU PSSM was subsequently used for searching AlgU binding sites in the genome of *A. vinelandii*. Predicted AlgU biding sites with a score ≥ 0.9 were found in the regulatory region of 117 genes, a number like that found for *P. aeruginosa*. 420 genes showed a predicted AlgU-dependent promoter with a score of 0.81 (S10 Table).

For comparative purposes, the prediction of potential AlgU promoters was extended for the *P. fluorescens* SBW25 and *A. chroococcum* Ac-8003 genomes (S11 and S12 Tables). The size of the predicted regulons was conserved, with about 100 motifs with a score ≥ 0.9. Interestingly, the genes encoding an OsmC-like protein, a peptidyl-prolyl cis-trans isomerase, a transaldolase TalB, the RpoH sigma factor, or genes required for trehalose synthesis or alginate production, were among the many genes conserved in the four bacteria (S13 Table). Of note, some of them were previously shown to be part of the AlgU regulon [77]. A sequence *logo* for the AlgU promoters identified in each bacterium was generated and as expected, it reflected the original promoter sequences used as input to develop the PSSM (Fig 4 and S4 Fig).

**Fig 4.**
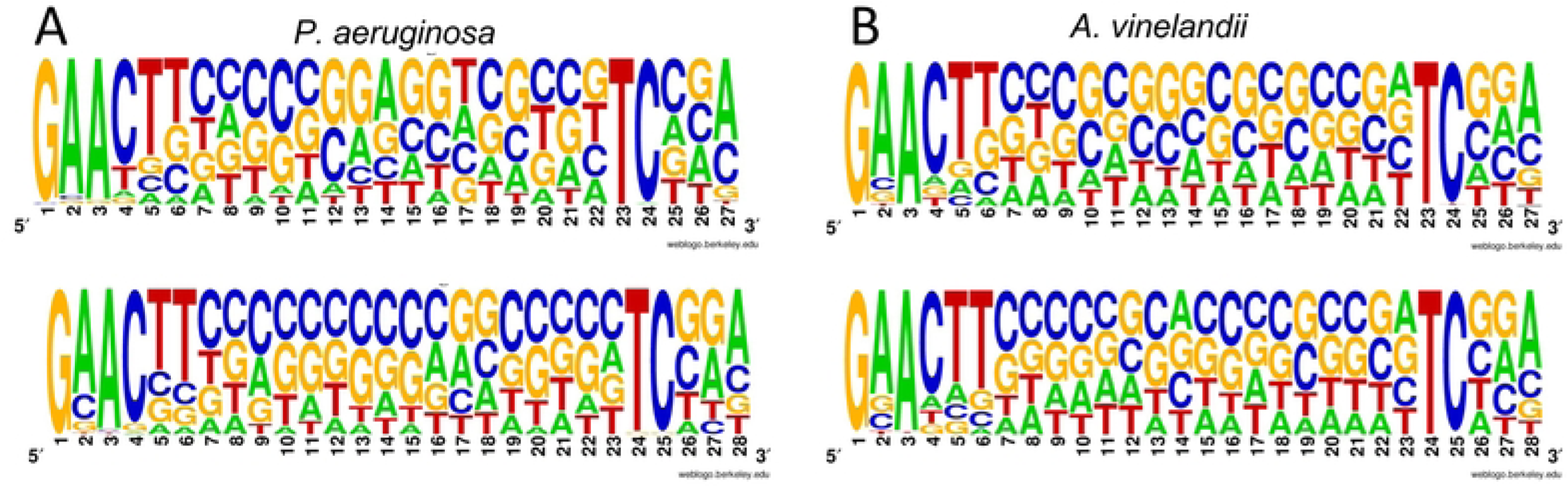
Sequence logo for AlgU DNA binding motifs. A PSSM was used to identify AlgU binding motifs in the genome of *P. aeruginosa* (A) or *A. vinelandii* (B) (see Materials and Methods section for details). The predicted AlgU binding motifs are shown with 16 (upper panels) or 17 (lower panels) bp spacers, between the −10 and −35 boxes. The relative sizes of the letters represent their frequency in the sequences. A comparative analysis between the predicted AlgU targets in the genome of *A. vinelandii* (S10 Table) and the gene products identified by our proteomic approach, under the positive control of this sigma factor (S4 and S7 Tables), allowed us to detect genes likely regulated by AlgU in a direct way. A total of 47 genes showed AlgU binding motifs in their regulatory region, while 53 genes were present in poly-cistronic operons with predicted AlgU promoters (S4 and S7 Tables, last three columns; S10 Table, fourth column). The presence of AlgU binding motifs for ten of these genes was shared among the *Azotobacter* and *Pseudomonas* species analyzed (highlighted in green, S13 Table), while for 20 genes it was restricted to *A. vinelandii* and *A. chroococcum* (highlighted in blue, Table S13). As expected, these latter set of genes, include those involved in ARs production and nitrogen fixation, but also include genes encoding an ATP synthase (Avin_52150-Avin_52210), an alcohol dehydrogenase (Avin_41760) or phosphoenolpyruvate carboxykinase (Avin_05450), among others. The importance of these genes in the physiology of the *Azotobacter* species deserves future investigations.

## Concluding remarks

In summary, we have reported the AlgU regulon during both, vegetative and encysting conditions in *A. vinelandii*. Although we detected molecular targets that explained processes previously documented under the control of AlgU (such as flagella biogenesis, alginate production or oxidative stress resistance), this work further expands our knowledge about the function of this sigma factor in *A. vinelandii.* AlgU is required for losing the flagella during the early steps of differentiation and agrees with the increase in the activity of this sigma factor (Fig 1). However, our data indicate that AlgU is also needed for the metabolic switch that takes place upon encystment. The *algU* mutant could not to produce ARs (Fig 3), implying that the replacement of the phospholipids of the cell membrane does not occur. A total of 337 proteins were found under the positive control of AlgU (S4 and S7 Table). Among these, the corresponding genes of 100 proteins showed predicted AlgU promoters in their regulatory region. The presence of AlgU binding motifs for some orthologous of *P. aeruginosa* or *P. fluorescens* was conserved but others were exclusive of *A. chroococcum* and *A. vinelandii* suggesting that the AlgU regulon is flexible and optimized for each bacterium.

## Acknowledgments

This work was supported by a grant from Programa de Apoyo a Proyectos de Investigación e Innovación Tecnológica, UNAM (PAPIIT) IN209521 to Cinthia Núñez. We thank J. Guzmán for her technical support and E. Bustos and J. Yañez for oligonucleotide synthesis and DNA sequencing services. SC-P was a recipient of a DGAPA, UNAM postdoctoral fellowship.

## Author contributions

**Conceptualization**: Sangita Chowdhury-Paul, Victoria Pando-Robles and Cinthia Núñez

**Formal analysis**: Sangita Chowdhury-Paul, Iliana C. Martínez-Ortíz, Enrique Merino and Cinthia Núñez

**Funding acquisition**: Guadalupe Espín and Cinthia Núñez

**Investigation**: Sangita Chowdhury-Paul, Enrique Merino and Cinthia Núñez **Methodology:** Sangita Chowdhury-Paul, Iliana C. Martínez-Ortiz, Soledad Moreno and Enrique Merino

**Project Administration:** Cinthia Núñez

**Resources:** Cinthia Núñez and Guadalupe Espín

**Supervision:** Sangita Chowdhury-Paul, Iliana C. Martínez-Ortiz, Victoria Pando-Robles, Soledad Moreno and Enrique Merino

**Validation:** Iliana C. Martínez-Ortiz, Victoria Pando-Robles, Enrique Merino and Cinthia Núñez.

**Writing-original draft:** Sangita Chowdhury-Paul, Enrique Merino and Cinthia Núñez

**Writing-review & editing:** Iliana C. Martínez-Ortiz, Victoria Pando-Robles, Guadalupe Espín, Enrique Merino and Cinthia Núñez

## Supporting Information

**S1 Fig.** Visualization of protein-protein interaction network by String 10.0 using the 126 proteins with altered expression in the absence of the sigma factor AlgU, during vegetative growing conditions. Interaction nodes such as those constituted by proteins involved in lipids metabolism (green circle), central metabolism (blue circle), flagella biogenesis and motility (cyan circle), trehalose synthesis (black circle) and enzymes for alginate production (pink circle) are indicated. Disconnected nodes are hided; the network was generated using an interaction score of 0.7.

**S2 Fig.** Visualization of protein-protein interaction network generated by String 10.0 from the 305 down-represented proteins in the absence of the sigma factor AlgU, during encysting conditions. Interaction nodes such as those constituted by proteins involved in ribosome assembly (red circle), nitrogen fixation (blue circle), amino acid metabolism (cyan circle), respiration (black circle), central metabolism (green circle) and enzymes for alginate (pink circle) or trehalose (yellow circle) production are indicated. Disconnected nodes are hided; the network was generated using an interaction score of 0.7.

**S3 Fig.** Visualization of protein-protein interaction network by String 10.0 from the 184 upregulated proteins in the absence of the sigma factor AlgU, during encysting conditions. Interaction nodes such as those constituted by proteins involved in amino acids (green circle), aminoacyl-tRNA (cyan circle), or fatty acid (black circle) biosynthesis are indicated. Disconnected nodes are hided; the network was generated using an interaction score of 0.7.

**S4 Fig. AlgU sigma factor binding motifs.** AlgU DNA binding motifs. A Position-Specific Scoring Matrix was used to identify AlgU binding motifs in the genome of *P. fluorescens* (A) or *A. chroococcum* (B) (see Materials and Methods section for details). The predicted AlgU binding motifs with are shown with 16 (upper panels) or 17 (lower panels) bp spacers between the −10 and −35 boxes.

**S1 Table.** DNA sequences of the primer pairs used in the present work.

**S2 Table.** List of experimentally determined AlgU promoters of *P. aeruginosa* that served as a reference for generating the Position-Specific Scoring Matrix.

**S3 Table.** Total proteins identified in the *algU* mutant under vegetative growth as compared to wild-type strain.

**S4 Table.** List of proteins downregulated in the absence of AlgU as compared to the wild-type strain, under vegetative growing conditions.

**S5 Table.** List of proteins upregulated in the absence of AlgU as compared to the wild-type strain, under vegetative growing conditions.

**S6 Table.** Total proteins identified in the *algU* mutant under encystment-induced conditions as compared to the wild-type strain.

**S7 Table.** List of proteins downregulated in the absence of AlgU as compared to the wild-type strain, under encystment-induced conditions.

**S8 Table.** List of proteins upregulated in the absence of AlgU as compared to the wild-type strain, under encystment-induced conditions.

**S9 Table.** Predicted AlgU-dependent promoters in the PAO1 genome.

**S10 Table**. Predicted AlgU-dependent promoters in the *A. vinelandii* DJ genome.

**S11 Table.** Predicted AlgU-dependent promoters in the *P. fluorescens* SBW25 genome.

**S12 Table.** Predicted AlgU-dependent promoters in the *A. chroococcum* Ac-8003 genome.

**S13 Table.** Comparative analysis of the predicted AlgU-dependent promoters in *A. vinelandii* with their orthologous in *A. chroococcum, P. aeruginosa* and *P. fluorescens*.

